# *Xenopus laevis* lack the critical sperm factor PLCζ

**DOI:** 10.1101/2023.02.02.526858

**Authors:** Rachel E. Bainbridge, Joel C. Rosenbaum, Paushaly Sau, Anne E. Carlson

## Abstract

Fertilization of eggs from the African clawed frog *Xenopus laevis* is characterized by an increase in cytosolic calcium, a phenomenon that is also observed in other vertebrates such as mammals and birds. During fertilization in mammals and birds, the transfer of the soluble PLCζ from sperm into the egg is thought to trigger the release of calcium from the endoplasmic reticulum (ER). Injecting sperm extracts into eggs reproduces this effect, reinforcing the hypothesis that a sperm factor is responsible for calcium release and egg activation. Remarkably, this occurs even when sperm extracts from *X. laevis* are injected into mouse eggs, suggesting that mammals and *X. laevis* share a sperm factor. However, *X. laevis* lacks an annotated *PLCZ1* gene, which encodes the PLCζ enzyme. In this study, we attempted to determine whether sperm from *X. laevis* express an unannotated *PLCZ1* ortholog. We identified *PLCZ1* orthologs in 11 amphibian species, including 5 that had not been previously characterized, but did not find any in either *X. laevis* or the closely related *Xenopus tropicalis*. Additionally, we performed RNA sequencing on testes obtained from adult *X. laevis* males and did not identify potential PLCZ1 orthologs in our dataset or in previously collected ones. These findings suggest that PLCZ1 may have been lost in the *Xenopus* lineage and raise the question of how fertilization triggers calcium release and egg activation in these species.

## Background

For most animals, fertilization triggers a rapid increase in the calcium of the egg cytoplasm. This elevated calcium then initiates polyspermy blocks, which prevent multiple sperm from entering an already fertilized egg, and embryonic development (Denninger et al., 2014; Swann and Lai, 2016). The sperm-derived enzyme phospholipase C zeta (PLCζ) is proposed to signal this process in mammals (Nomikos et al., 2017; Saunders et al., 2002) and birds (Coward et al., 2005) by increasing the egg inositol trisphosphate (IP_3_). This increase in IP_3_ then triggers calcium release from the endoplasmic reticulum (ER) (Bedford-Guaus et al., 2011; Cox et al., 2002; Ito et al., 2008; Mizushima et al., 2014; Mizushima et al., 2007; Ross et al., 2008; Saunders et al., 2002; Yoneda et al., 2006). It is not yet known whether other animals use PLCζ in the same way.

We sought to determine if the African clawed frog (*Xenopus laevis*) uses PLCζ in fertilization. Fertilization of *X. laevis* eggs increases cytoplasmic calcium (Grey et al., 1982; Kline, 1988), which then activates a chloride channel called TMEM16A to depolarize the membrane and initiate the fast block to polyspermy (Wozniak et al., 2018a). The initiation of the fast block to polyspermy requires PLC activation (Wozniak et al., 2018b), and we previously explored a role for egg PLCs in signaling the fast block. *X. laevis* eggs express three PLC isoforms (Plcg1, Plcb1, and Plcb3) (Komondor et al., 2022). However, we found that the signaling pathways that typically activate these isoforms are not involved in the fast block (Komondor et al., 2022). We therefore considered the possibility that *X. laevis* sperm donates PLCζ to the egg, as occurs in mammals and birds. The observation that injecting extracts from *X. laevis* sperm into mouse eggs leads to an elevation of cytoplasmic calcium comparable to fertilization in mouse eggs supports the idea that PLCζ may be involved in fertilization in *X. laevis* (Dong et al., 2000). However, no PLCζ-encoding gene (*PLCZ1*) has been identified in the genomes of *X. laevis* or the closely related species *X. tropicalis*.

## Results

At the outset of our study, we conducted a search for PLCζ in amphibians. The class Amphibia is comprised of three orders: tailless frogs and toads (Anura), newts and salamanders (Urodela), and limbless caecilians (Gymnophiona). Using blastp with mouse *Plcz1* as our query against the non-redundant sequence database (nr), we identified *PLCZ1* orthologs in six unique amphibian species: the Gaboon caecilian (*Geotrypetes seraphini*), two-lined caecilian (*Rhinatrema bivittatum*), common frog (*Rana temporaria*), Tibetan frog (*Nanorana parkeri*), common toad (*Bufo bufo*), and the Asiatic toad (*Bufo gargarizans*) (Fig. 1a). These putative *PLCZ1* orthologs, along with other PLC isoforms found in the same species, were included in an alignment with PLCs from representative species including mammals, birds, reptiles, and lungfish, which served as an outgroup (Fig. 1b). The annotated *PLCZ1* orthologs were found to cluster with other PLCZ1 isoforms in the alignment (Fig. 1c), supporting their classification as PLCζ isozymes.

**Figure 1.**
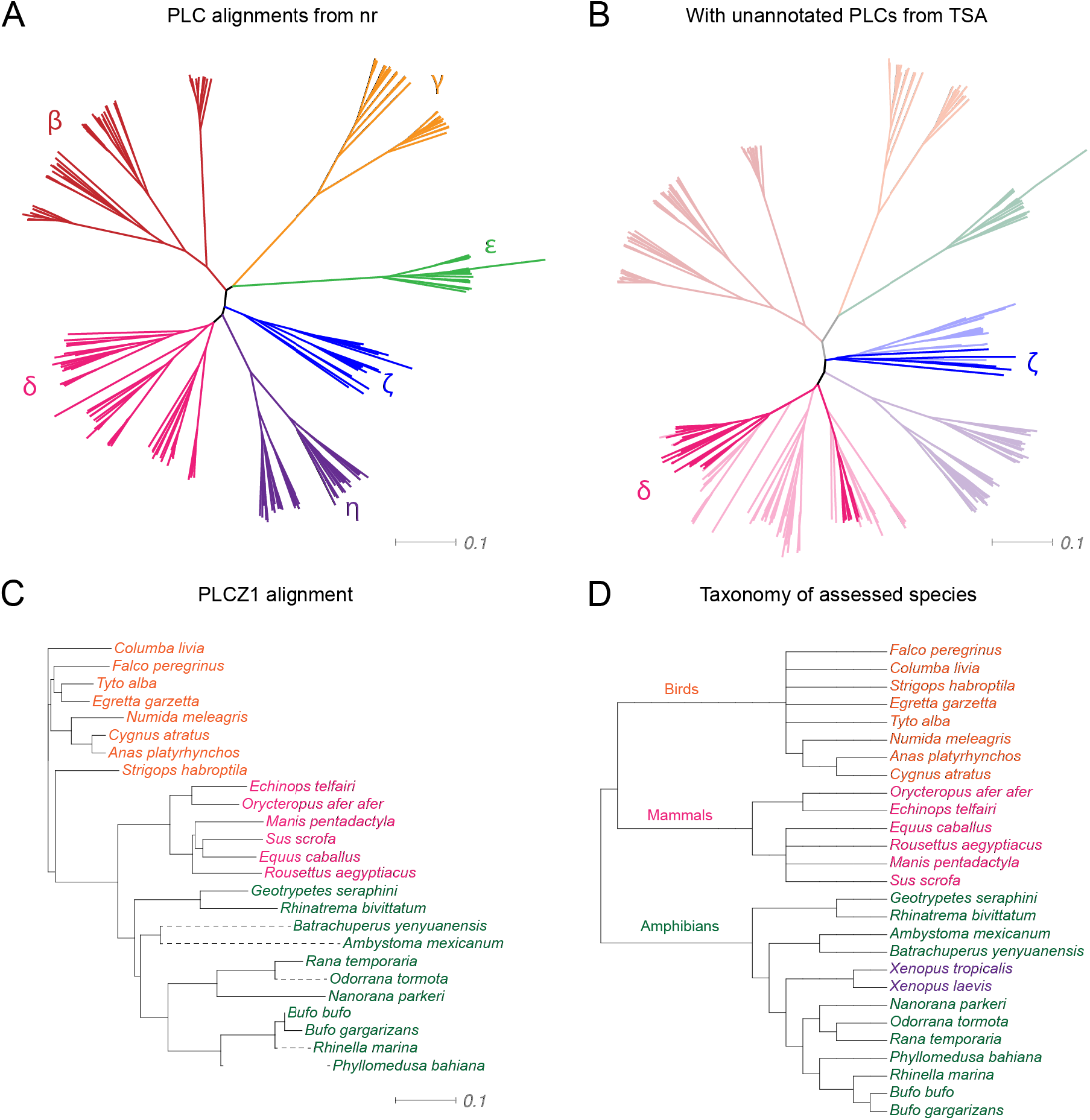
Identification of amphibian PLCζ isozymes. **a** Phylogram displaying the alignment of annotated PLC isoforms from representative mammals, birds, and amphibians. **b** Alignment of unannotated *PLCZ1* homologs from the amphibian transcriptome shotgun assembly (TSA) dataset. Reference sequences from (**a**) are included. **c** Phylogram representing the alignment of *PLCZ1* orthologs (branches indicated with dashed lines) identified in (**b**) with annotated sequences from the non-redundant nucleotide database (solid branches). **d** Cladogram of species from (**c**); *Xenopus* species (for which no *PLCZ1* orthologs could be identified) are indicated in purple.

To expand our search for PLCZ1 isozymes in amphibians, we used tblastn with Plcz1 from the Tibetan frog (*Nanorana parkeri*) as our query against the TSA database. This analysis identified both *PLCZ1* and *PLCD1-4* orthologs. We included these sequences in an alignment with PLCs from a variety of representative species, which clearly identified five additional *PLCZ1* orthologs: the Yenyuan stream salamander (*Batrachuperus yenyuanensis*), axolotl (*Ambystoma mexicanum*), concave-eared torrent frog (*Odorrana tormota*), cane toad (*Rhinella marina*), and the neotropical leaf-frog (*Phyllomedusa bahiana*) (Fig. 1b). These additional sequences also grouped with other PLCZ1 orthologs from amphibian species in a way that matched the taxonomic classification of the assessed species (as shown in Figs. 1c&d).

We conducted transcriptomic analysis to determine if an unannotated *PLCZ1* ortholog is expressed in *X. laevis*. In mammals, the PLCζ protein is exclusively found in sperm (Session et al., 2016), but mature sperm are transcriptionally quiescent (Kierszenbaum and Tres, 1975) and not suitable for RNA-seq. We therefore performed RNA-seq on whole testes, recognizing that not all RNA present in the testes represents proteins found in sperm, but RNA that encode sperm proteins should be present in the testes. We obtained RNA from the testes of three individual adult male *X. laevis* frogs and performed RNA-seq on these three samples. Of the contiguous RNA-seq reads, 48-51% were aligned to the *X. laevis* genome. We observed at least one transcript in one testis from an individual frog for 33,377 genes. We performed gene ontology (GO) analysis on 6,000 of the most highly expressed genes among these testis-derived transcripts. Of these, 48.3% were identified with GO biological process terms, many of which were important for general cell maintenance (*e.g*. translation, protein folding, cell cycle). Some were also involved in specialized processes such as spermatogenesis. Our overall results were highly correlated (Fig. S1) and generally matched related datasets (Fig. 2). In our dataset, we identified several essential sperm transcripts, including deleted in azoospermia-like (*dazl*) (Houston and King, 2000) and the testes-specific histone (*h1-10*) (Oikawa et al., 2020; Shechter et al., 2009). Other enriched transcripts, such as *astl2b, rflcii*, and *ribc1*, are known to be specifically expressed in *X. laevis* testis (Session et al., 2016) or as proteins in sperm (Teperek et al., 2014). Some of these transcripts, including *rflcii, cfh*, and *tubb4b*, have also been shown to be required for fertility in other species (Table 1) (Agarwal et al., 2016; Ferlin et al., 2012; Maccarinelli et al., 2017; Sakaue et al., 2010).

**Figure 2.**
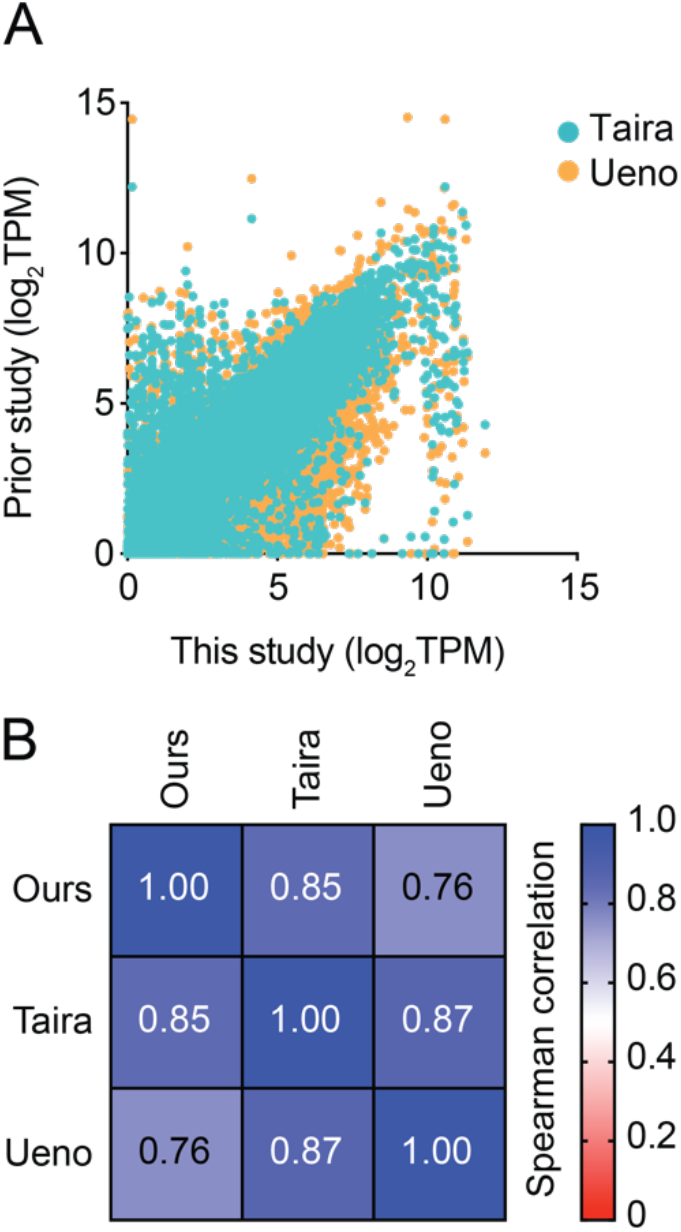
Comparison of our RNA-seq data with two other published datasets. **a** Plot showing transcripts per million (TPM) for matched transcripts between this study (X-axis) and two datasets in a previous study (Session et al., 2016). **b** Array representing the Spearman rank correlation between each dataset in (**a**).

**Table 1.**
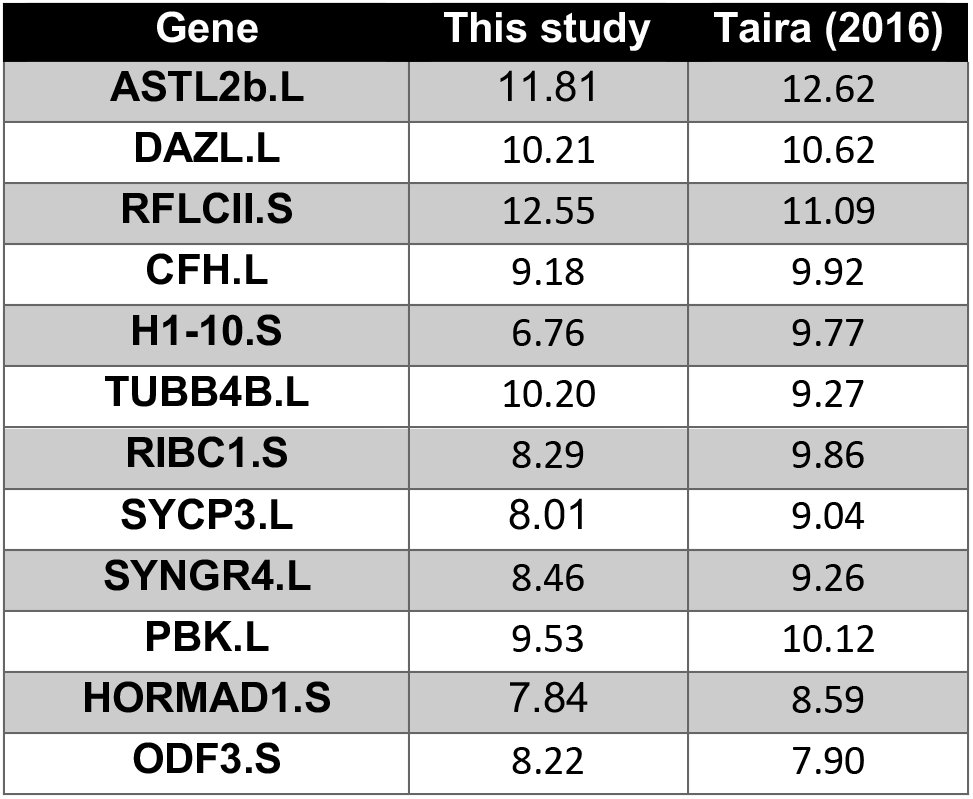
Comparative gene expression in testes of *X. laevis*. Expression of testis-associated genes between our averaged datasets and the Taira dataset (Session et al. 2016).

To identify sequences in the unmapped RNA that may belong to a *PLCZ1* gene, we ran a blastx search of the reads that hadn’t been matched to any known genes against a database of annotated *PLCZ1* sequences from mammals and amphibians. This search identified 360 reads that had a very low E-value (less than 10^-6^) and covered a specific region (residues 170-654) of a representative PLCZ1 protein sequence from the neotropical leaf frog, *Phyllomedusa bahiana*. However, when we used a blastn search to compare these reads to the amphibian nr database, we discovered that most of them matched to *PLCD4* or other isoforms (Supporting files). This makes sense in that PLCζ is the only PLC isoform that lacks a pleckstrin homology (PH) domain but is otherwise like PLCδ (Fig. 3). Our results suggest that the unmapped reads in our dataset arose from incomplete contigs of annotated genes.

**Figure 3.**
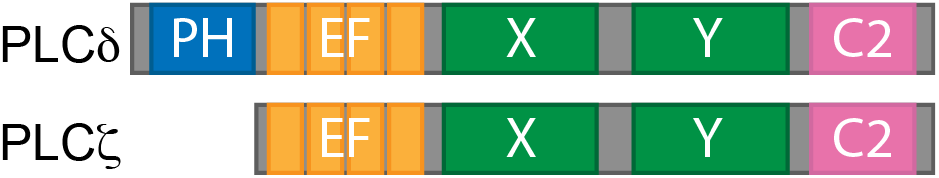
Domain architecture of PLC isoforms PLCδ and PLCζ. Schematic representation of the domain structures: pleckstin homology (PH) domain, EF hands, X and Y catalytic domains, and the C-terminal C2 domain.

We searched the RNA-seq dataset for transcripts encoding PLC isoforms and found transcripts for PLCβ, PLCδ, and PLCγ isoforms (Fig. 4a). Of these, the *plcd4* transcript was the most abundant. *plcd4* was also the most abundant PLC transcript in testis RNA-seq datasets from the inbred *X. laevis* J strain (Session et al., 2016) and the *X. tropicalis* Nigerian strain, both of which lacked *PLCZ1*. By comparison, testes from the Oriental fire-bellied toad *Bombina orientalis* contained *plcz1* transcripts but had substantially reduced levels of *plcd4* compared to *Xenopus* (Fig. 4a). By comparison, data obtained from adult (6 months old) mouse testis revealed that *Plcz1* was the most abundant transcript, followed by *Plcd4* (Fig. 4b) (Huang et al., 2021).

**Figure 4.**
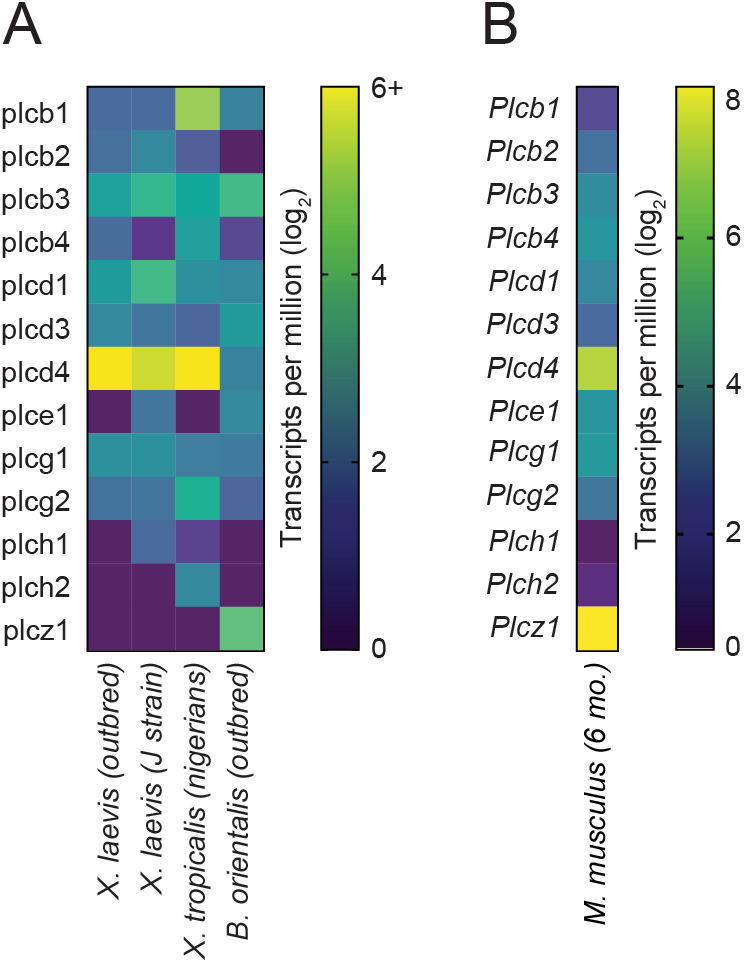
Heatmap of PLC expression in anuran and mouse testis. **a** Heatmaps of the expression levels (shown as log_2_ transformed transcripts per million) of annotated PLC genes from the indicated animals. The *X. laevis* (outbred) data obtained here, *X. laevis* J strain from GSE73419 (Session et al., 2016), *X. tropicalis* from GSM5230669 (unpublished), and *B. orientalis* from GSE163874 (unpublished). **b** Expression levels of PLC isoforms in the testis of adult (6 month) mice (GSE181426) (Huang et al., 2021).

## Discussion

As in other species, fertilization in *X. laevis* triggers an elevation of cytosolic calcium in the egg (Runft et al., 2002). This calcium is released from the ER by a pathway that requires PLC (Runft et al., 1999). Several mechanisms have been proposed to explain the trigger that initiates this process. In mammals and birds, the release of PLCζ from the sperm into the egg during fertilization is thought to trigger the increase in calcium in the egg cytoplasm (Swann and Lai, 2016).

Because of the role that it plays in mammalian and avian fertilization, we were surprised to find that there was no annotated ortholog of *PLCZ1* in *X. laevis*. This prompted us to investigate whether other amphibians possess this enzyme. In our search, we used both nucleotide and transcriptome shotgun assembly (TSA) databases, using an amphibian *PLCZ1* ortholog as a query to increase our chances of finding a match. We were ultimately able to identify *PLCZ1* orthologs in 11 amphibian species (Fig. 1a-b). Although this represents a small portion of the amphibian transcriptomes available in the TSA database, the lack of *PLCZ1* in the TSA search does not necessarily mean that it is not expressed in the associated species, as many amphibian transcriptome datasets do not include testes, where PLCζ is known to be expressed.

We aligned the amino acid sequences of the PLCZ1 protein in amphibians to those of the same protein in other animals, as well as to the sequences of other PLC subtypes, including the beta, gamma, delta, epsilon, and eta isoforms. This comparison included species with PLCζ enzymes with a verified role in fertilization, such as mammals and birds. By aligning the sequences of all PLC isoforms from each species, we found that the isoforms tended to group together primarily by subtype (Fig. 1a-b) and within subtype by order (Fig. 1c). This pattern of grouping by order is also reflected in the taxonomic classification of the species we examined (Fig. 1d). With the exceptions of *X. laevis* and *X. tropicalis*, every amphibian in our dataset transcribed a *PLCZ1* ortholog. These results suggest that *PLCZ1* may have been lost in the *Xenopus* lineage.

## Conclusions

Since the *Xenopus* species examined in our study lack a PLCζ isozyme, how fertilization activates PLC to trigger calcium release from the ER in these frogs is yet to be determined. Several sperm factors other than PLCζ have been proposed to initiate this process, including citrate synthase (Harada et al., 2011; Harada et al., 2007) and post-acrosomal WWP-domain-binding protein (PAWP) (Wu et al., 2007). For *Xenopus*, it’s possible that a sperm derived protein does not signal an increase of the cytoplasmic calcium in the egg. For example, another possibility is the “receptor model” of egg activation, in which a signaling cascade is activated by the binding of a sperm surface ligand to an egg receptor (Jaffe, 1990). *X. laevis* eggs express a PLCg isoform (Plcg1), which can be activated by phosphorylation of a critical residue in the active site (Kadamur and Ross, 2013), as well as PLCβ isoforms (Plcb1 and Plcb3), which are activated by GPCRs (Rhee, 2001; Smrcka et al., 1991). However, fertilization in *X. laevis* does not cause observable phosphorylation of the critical tyrosine residue of Plcg1, and inhibitors of either PLCγ or PLCβ pathway had no effect on fertilization (Komondor et al., 2022). Another alternative explanation is the “calcium bomb” hypothesis, which proposes that a burst of calcium is introduced to the egg by the sperm itself, resulting in egg activation and calcium release from the ER (Jaffe, 1983). In other model systems, introducing calcium alone was not sufficient to produce calcium release, indicating that other factors are required (Swann and Ozil, 1994). Whether calcium alone is sufficient to initiate the polyspermy blocks and embryonic development in *Xenopus* eggs is yet to be determinized. Further research is needed to fully understand the mechanisms behind activation of *Xenopus* eggs.

## Methods

### Retrieval of amphibian PLCZ1 sequences

We searched the BLAST non-redundant protein sequence database for amphibian *PLCZ1* orthologs using the protein sequence of mouse Plcz1 as a query. We retrieved annotated orthologs from six amphibian species and included them, along with sequences encoding other PLC isoforms (beta, gamma, delta, epsilon, and eta) in an alignment of representative PLC sequences from amphibians, birds, and mammals using Kalign (Lassmann, 2020). We used the neighbor-joining phylogenetic output from this alignment to generate an unrooted tree with Dendroscope (Huson and Scornavacca, 2012).

We expanded our search for *PLCZ1* orthologs by performing a reverse translated tblastn search using the sequence of the *Nanorana parkerii PLCZ1* as a query against the amphibian transcriptome shotgun assembly (TSA) database. We retrieved more than 100 sequences of possible *PLCZ1* orthologs, which we translated using getorf (EMBOSS), selecting the longest open reading frames from each transcript. We aligned these sequences with the representative PLC sequences that we had previously obtained from amphibians, birds, and mammals to distinguish genuine *PLCZ1* orthologs from hits that encoded other PLC subtypes. Through this analysis, we identified genes encoding *PLCZ1* in 5 additional amphibian species. Finally, we used InterProScan (Nomikos et al., 2005) to validate the potential PLCZ1 orthologs that we had identified by looking for the absence of a pleckstrin homology (PH) domain, which is a defining feature of PLCζ enzymes.

### Animals

All animal procedures were conducted using acceptable standards of humane animal care and approved by the Institutional Animal Care and Use Committee (IACUC) at the University of Pittsburgh. *X. laevis* adults were obtained commercially (Nasco) and kept in a controlled environment with a 12-hour light/dark cycle at 20°C.

### RNA isolation

To obtain RNA, we euthanized sexually mature *X. laevis* males by immersing them in a solution of tricaine-S (3.6 g/L, pH 7.4) for 30 minutes. We then removed the testes and carefully cleaned them to remove fat and vascular tissue. We prepared tissue for RNA isolation by freezing it with liquid nitrogen and grinding it into a powder with a mortar and pestle. We then isolated RNA following instructions provided by RNeasy and QIAshreddder kits (QIAgen).

### RNA-seq library preparation and data acquisition

Before preparing the RNA-seq library, we first determined the integrity of the RNA using electrophoresis and measured its concentration by Qubit (Life Technologies). We then used Illumina TruSeq mRNA kit (Illumina) with modified protocol: SuperScript IV (Invitrogen) was used for first strand synthesis, the library amplified with 10 cycles of PCR, and the amplified library was cleaned up with 35 μL AMPureXP beads (Beckman Coulter). The library was sequenced with 75 bp paired-end mRNA reads on an Illumina NextSeq500 platform with a Mid Output 150 flowcell (Illumina).

### RNA-seq analysis

Sequencing reads were uploaded to the public server at usegalaxy.org (Afgan et al., 2018). The reads were then aligned to *Xenopus laevis* genome (version 10.1) using HISAT2 (Galaxy version 2.2.1+galaxy0) with default settings for paired reads. Next, aligned fragments were mapped and quantified with featureCounts (Galaxy version 2.0.1+galaxy2) (Liao et al., 2014) using the Xenbase gene model as a reference.

### BLAST search for unmapped PLCZ1

Command line BLASTx was used to search unmapped RNA-seq reads against a database containing annotated PLCZ1 protein sequences from *Mus musculus* (AAI06769.1), *Homo sapiens* (AAI25068.1), *Bufo gararizans* (XP_044137439.1)*, Bufo bufo* (XP_040291278.1)*, Rhinatrema bivittatum* (XP_029453199.1), and *Nanorana parkeri* (XP_018417886.1). We used an E value cutoff of 10^-6^, which produced 360 reads. We then aligned these reads to the nucleotide sequence of *PLCZ1* from *Phyllomedusa bahiana*, which produced a noncontiguous alignment covering residues 170-654 of *PLCZ1*. To verify that these reads were not attributable to other PLC isoforms, we performed a multiple query BLAST search with these reads against amphibian sequences from the non-redundant nucleotide (nt) database.

### Comparative transcriptome analysis

To compare transcript levels, we took the average of three individual experiments and reported transcript abundance as the number of transcripts per million. We deposited this data with the NCBI Gene Expression Omnibus (Edgar et al., 2002), which can be accessed using accession number GSE224304. To ensure that our data could be compared to previously existing data, we matched the transcriptome of our dataset against a GEO dataset for *X. laevis* (J strain) (GSE73419) (Session et al., 2016). We identified 13,277 genes to use for direct comparison, which we then ranked and compared using Spearman’s rank correlation in Prism (Graphpad).

To compare the expression of PLC subtypes in the testes of different organisms, we obtained GEO data for *Xenopus laevis* (J strain) (GSE73419), *Xenopus tropicalis* (nigerians) (GSM5230669), *Bombina orientalis* (GSE163874), and *Mus musculus* (GSE181426). We processed the expression data using a log_2_ transformation and created a heatmap using Prism (Graphpad) to visualize the interspecies comparison.

## List of abbreviations

Abbreviations used are:

PLC: phospholipase C
IP_3_: inositol triphosphate
ER: endoplasmic reticulum
TMEM16A: TransMEMbrane 16A
TSA: transcriptome shotgun assembly
nr: non-redundant sequence database
RNA-seq: RNA sequencing

## Availability of data and materials

The datasets generated and/or analysed during the current study are available in the Gene Expression Omnibus, which can be accessed using accession number GSE224304.

## Author Contributions

Conceptualization: REB, JCR, AEC

Data curation: REB, JCR, PS, AEC

Formal analysis: REB, JCR

Investigation: REB, JCR, AEC

Methodology: REB, JCR

Project administration: REB, JCR, AEC

Resources: REB, JCR, AEC

Visualization: REB, JCR, AEC

Writing – original draft: REB, JCR, AEC

Writing – reviewing & editing: REB, JCR, PS, AEC

## Acknowledgements

We thank D. Summerville for excellent technical assistance, and M.T. Lee for stimulating conversations.

---

**Figure S1.**
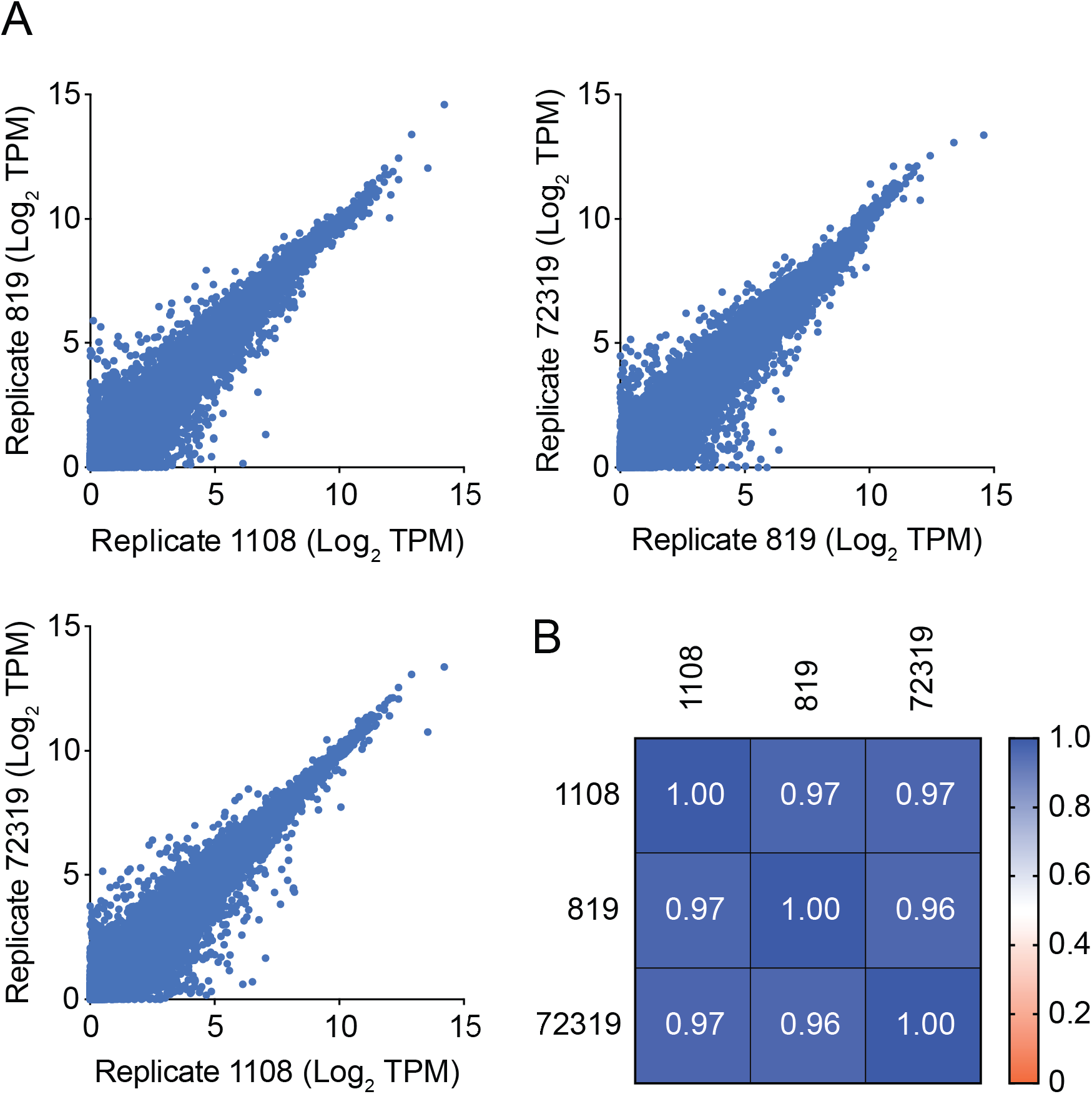
**a** Correlation in log_2_ transcripts per million for each gene across experimental replicates. **b** Array representing the Spearman rank correlation between each dataset in (**a**).

